# Stacked binding of a small molecule PET tracer to Alzheimer’s tau paired helical filaments

**DOI:** 10.1101/2022.09.30.510175

**Authors:** Gregory E. Merz, Matthew J. Chalkley, Sophia Tan, Eric Tse, Joanne Lee, Stanley B. Prusiner, Nick A. Paras, William F. DeGrado, Daniel R. Southworth

## Abstract

Neurodegenerative diseases (NDs) are characterized by the formation of amyloid filaments that adopt disease-specific conformations in the brain. Recently developed small molecules hold promise as diagnostics and possible therapeutics for NDs, but their binding mechanisms to amyloid filaments remain unknown. Here, we used cryo–electron microscopy (cryo-EM) to determine a 2.7 Å structure of Alzheimer’s disease patient-derived tau paired-helical filaments incubated with the GTP-1 PET probe. GTP-1 is bound stoichiometrically along an exposed cleft of each protofilament in a stacked arrangement that matches the fibril’s symmetry. Multiscale modeling revealed favorable pi-pi aromatic stacking interactions between GTP-1 molecules that, together with small molecule–protein contacts, result in high affinity binding. This binding mode offers new insight into designing compounds for diagnosis and treatment of specific NDs.

**One Sentence Summary:** Cryo-EM structure reveals a novel stacked arrangement of the GTP-1 PET ligand bound to Alzheimer’s disease tau filaments.

## Main text

The accumulation of misfolded tau proteins in the brain is a hallmark of the large subset of neurodegenerative diseases (NDs) known as tauopathies (*1, 2*), the most common and widely studied of which is Alzheimer’s disease (AD) (*3*). The spread of tau deposits, known as neurofibrillary tangles (NFTs) in AD, parallels neuronal loss and cognitive impairment (*4, 5*) and serves as a marker for disease progression (*6*). Moreover, accumulation of NFTs appears to be the final product of a process in which soluble tau misfolds into amyloid filaments that self-propagate and transmit as prions across neurons via synaptic junctions (*7*). Prions were first identified in the scrapie prion protein (PrP^Sc^), which causes Creutzfeldt-Jakob (CJD), Gerstmann-Sträussler-Scheinker (GSS), and other incurable diseases (*8, 9*) in which amyloids also accumulate with disease progression. Structures determined by cryo-EM of tau filaments purified from patient brains have revealed that tau filaments adopt different cross-β sheet conformations of the microtubule binding repeat region among different NDs (*10–15*). For example, AD filaments are comprised of 3R and 4R isoforms and adopt a C-shaped fold, while in Pick’s disease, 3R tau forms an elongated J-shaped fold, and in corticobasal degeneration (CBD), 4R tau adopts a 4-layered β-strand arrangement (*14*). These distinct structural conformations have opened up the possibility of binding small molecules to different tau filament conformers for disease-specific targeting; here, we determined a cryo-EM structure of a small molecule bound to tau that reveals a potential mechanism for achieving site-specificity.

Small molecules that can discriminate among amyloid filaments (*16, 17*), and even strains of the same prions (*18, 19*), have been developed. However, the basis of this specificity is unknown. Despite this limitation, a number of promising tau-selective positron-emission tomography (PET) ligands for AD have been developed and tested *in vivo* (*20*). Many such molecules contain heterocyclic aromatic moieties, including Tauvid, a first-generation tau PET ligand that is FDA-approved and clinically available (*21*). While second-generation PET tracers have been developed to reduce off-target binding and optimize pharmacokinetic properties (*22, 23*), their direct binding to disease-relevant tau filament folds is undercharacterized. Docking studies have predicted that PET tracers bind end-to-end with the plane of the aromatic rings parallel to the fibril axis (*24, 25*), and a cryo-EM structure of the PET tracer APN-1607 at low-resolution (*26*) has been modeled to have the same orientation. On the other hand, cryo-EM studies of the small molecule Epigallocatechin gallate (EGCG, a compound known to disaggregate amyloid filaments *in vitro*) (*27*) bound to AD tau PHFs showed several unique densities. Model building indicated that the most well-defined of these densities represented EGCG molecules with their aromatic rings stacked perpendicular to the fibril axis; however, the molecular details of the interactions were not well resolved based on the density (*28*). Higher resolution co-structures are needed to identify the molecular features underlying site-specific binding modes of ligands to amyloid fibrils, and this knowledge will impact the design of conformationally specific diagnostic and therapeutic compounds.

Using cryo-EM, we sought to determine the co-structure of AD tau filaments and GTP-1 (<u>G</u>enentech <u>T</u>au <u>P</u>robe <u>1</u>), a high affinity (11 nM K_d_), second-generation tau PET tracer that is currently in clinical trials (Fig. 1A) (*29*). Tau filament samples were purified from the frontal cortex of a patient with AD as described previously (*10*) and showed high infectivity in a cell-based assay (*30*) (fig. S1). Samples were incubated with 20 μM GTP-1 prior to vitrification. The micrograph images and their 2D classification reveal well-resolved filaments primarily in the PHF conformation, with crossover distances ranging from 700–800 Å (fig. S2). A minor population of straight filaments (SFs) was also identified; however, further structural characterization was not feasible due to limited abundance (fig. S2). Using standard helical reconstruction methods (table S1 and Methods), a structure of the PHF was determined with an overall resolution of 2.7 Å (fig. S3) and is comprised of two protofilaments related by two-fold symmetry with a 2.37 Å rise and 179.45° twist (table S1), consistent with previously reported structures of PHFs prepared from AD brain tissue (*10, 12*). The central region surrounding the protofilament interface is at the highest resolution at ~2.5 Å and the periphery is at ~3.2 Å, indicating high resolution across the β-sheet core, as exhibited by well-resolved side chain densities (fig. S4A).

**Fig. 1.**
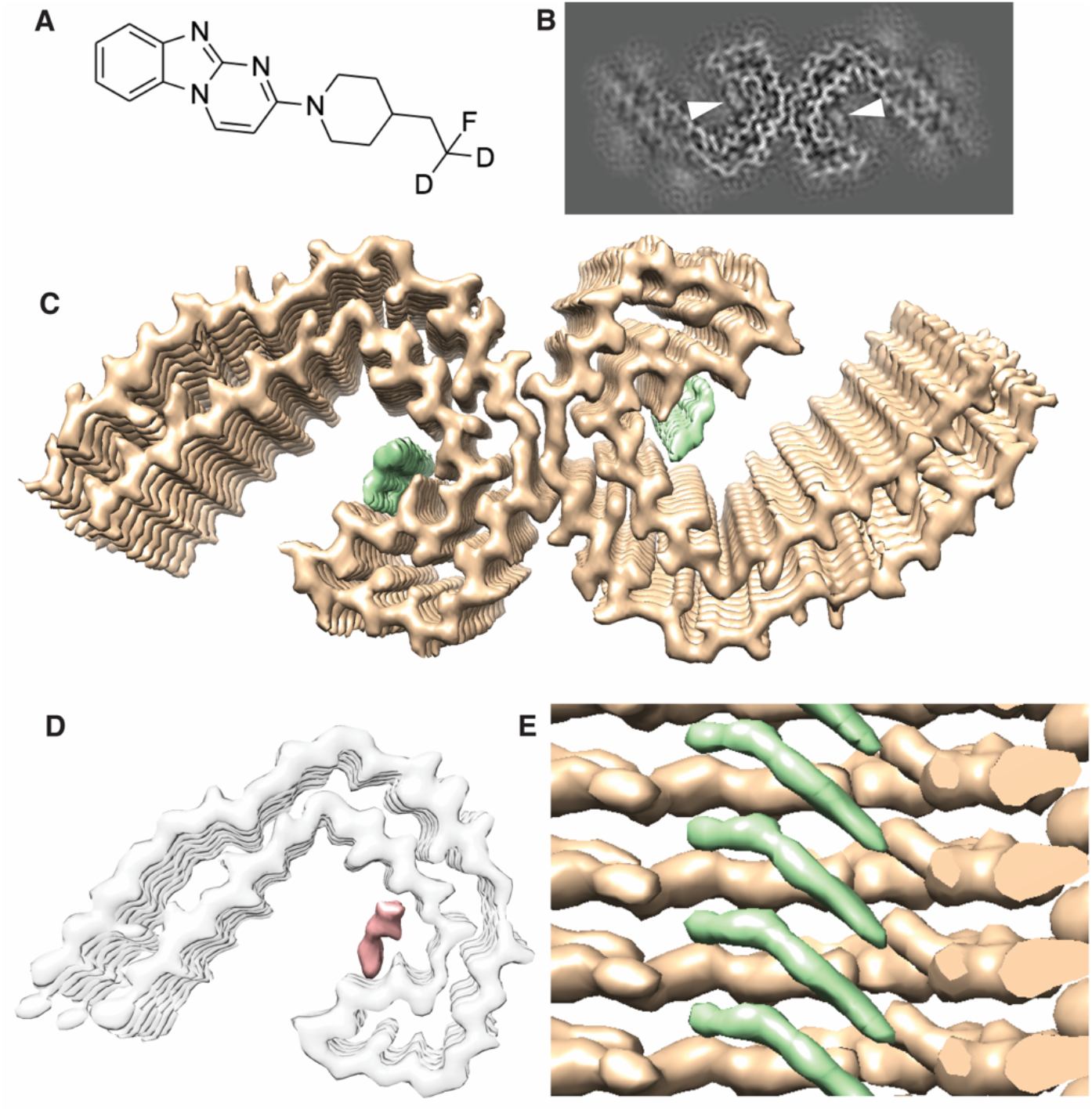
Cryo-EM map of AD tau PHF with density for bound GTP-1. (**A**) Chemical structure of GTP-1. (**B**) X-Y-slice view of the cryo-EM map of AD PHFs incubated with GTP-1. Extra density corresponding to GTP-1 is indicated by white triangles. (**C**) Cryo-EM map of tau PHF:GTP-1. Density corresponding to GTP-1 is colored in green. (**D**) Difference map (salmon density) between (C) and a previously determined apo-AD PHF map (EMDB: 0259), low-pass filtered to 3.5 Å. Density for the apo PHF protofilament (grey) is shown as a reference. (**E**) Side view of tau PHF:GTP-1 structure showing the ligand density (green) in a stacked arrangement with one molecule spanning across multiple rungs of the tau protofilament.

Remarkably, the structure reveals strong additional density that is indicative of the GTP-1 small molecule bound to a solvent-exposed cleft (Fig. 1, B and C); notably, this density appears identical in both protofilaments, indicating equivalent binding, considering two-fold symmetry was not enforced in the refinement. While other densities are present around the filament core, these are poorly resolved in comparison and similar to previously reported tau filament structures (figs. S4B and S5) (*10, 12*). Importantly, difference map analysis comparing the GTP-1 co-structure (tau PHF:GTP-1) to a previously determined PHF map (EMDB: 0259) (*12*) identifies that this density is uniquely present, with no additional density in the difference map (Fig. 1D). The lack of additional densities in our structure contrasts with earlier studies and indicates a highly specific interaction, as 20 μM GTP-1 is well above the measured IC50 (22 nM) (*29*). Notably, GTP-1 density indicates the compound is stacked in a geometric repeat that precisely matches that of protein monomers in the fibril (Fig. 1E). This arrangement contrasts previous studies reporting binding end to end or parallel to the fibril axis (*24–26*) but is similar to the stacked EGCG-tau model (*28*).

An atomic model of the tau portion of tau PHF:GTP-1 was achieved by docking and refinement of the previous PHF structure solved in the absence of exogenous ligand (Fig. 2A) (*12*). The protofilaments form the canonical C-shaped cross-β fold found in AD that is comprised of the 3R and 4R tau domains (residues 306–378) and interact laterally via the antiparallel PGGGQ motif (residues 332–336). The overall filament structure is nearly identical to previous structures of AD PHFs (α-carbon RMSD = 0.5 Å) (fig. S6A). GTP-1 is bound in the cleft comprised of residues 351–360 (Fig. 2A), adjacent to the three-strand β-helix (β5–7) in the protofilament. Small differences are seen in the sidechains of the residues lining the binding pocket, namely Gln 351, Lys353, Asp358, and Ile360 (fig. S6B).

**Fig. 2.**
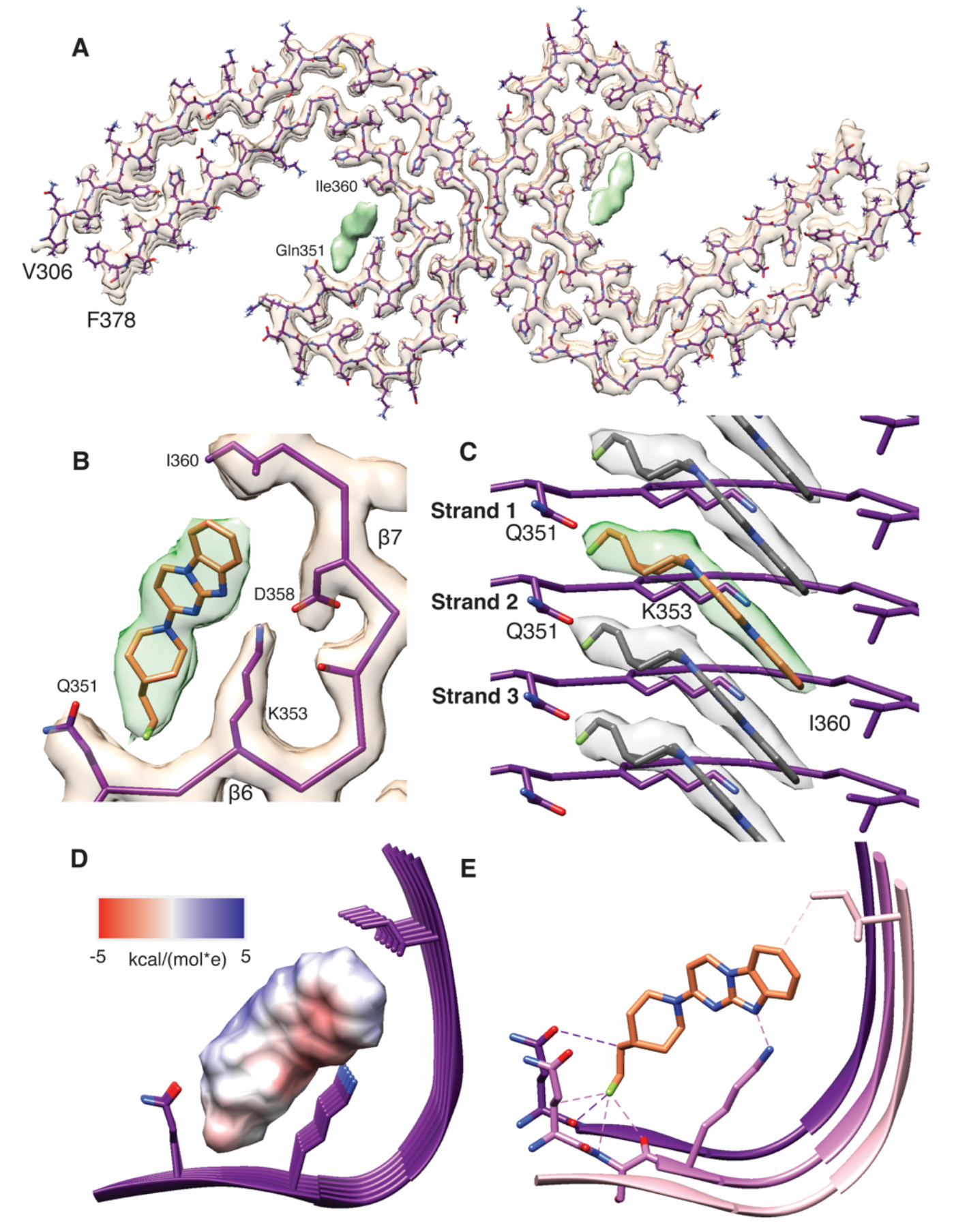
Atomic model of tau PHF and bound GTP-1. (**A**) Refined tau PHF atomic model fit into the PHF:GTP-1 density. (**B**) Map and model of the GTP-1 binding site with GTP-1 modeled into the density using a combination of molecule mechanics and DFT approaches. (**C**) Side view of tau PHF:GTP-1 model, showing individual GTP-1 molecules fit at an angle relative to the backbone and making contact across 3 rungs of tau. (**D**) GTP-1 electrostatic (Coulomb) potential surface representation showing complementarity to the GTP-1 binding pocket. (**E**) Close contacts (<3.5 Å) of GTP-1 with sites in the binding pocket.

Accurately modeling small molecule ligands is a notable challenge (*31*), and the tau PHF:GTP-1 structure presents additional difficulties due to the novel stacked arrangement of GTP-1 in which ligand-ligand interactions are likely making substantial contributions. Furthermore, while the tricyclic aromatic ring is rigid, the piperidine ring and fluoroethyl tail are highly flexible and difficult to model by standard methods (fig. S7). Our best modeling approach resulted from a combination of using molecular mechanics to generate conformers and then using density functional theory to perform constrained optimizations of dimers to capture small molecule–small molecule interactions, followed by final refinement with Phenix (*32*). The final modeled conformer yields excellent map-model agreement and is energetically reasonable (Fig. 2B, fig. S8, and table S2; see Methods). This map-model agreement, along with the fact that the ligand density has similar resolution to the adjacent filament structure (~2.6 Å) and remains present at high sigma threshold values, indicates near-complete occupancy (figs. S5B and S6).

GTP-1 binds in the C-shaped groove of the PHF filament comprised of strands β6 and β7, which are separated by a kink at Gly355 that creates a concave cleft complimenting the convex shape of the GTP-1 stack (Fig. 2B). We identify that each molecule of GTP-1 binds across three β-strands, making direct contacts with Gln351 in strand 1, Gln351 and Lys353 in strand 2, and Ile360 in strand 3, as well as the backbone between Gln351 and Lys353 in strands 1 and 2 (Fig. 2C). Notably, the piperidine ring and fluoroethyl tail of GTP-1 are parallel to the filament and project across two β-strands, making contact with sidechains and backbone atoms of Gln351 in both strands. Although the site is comprised of primarily polar residues, there is precise matching between the apolar portions of their sidechains and the apolar portions of the small molecule (Fig. 2D and fig. S9). The aliphatic carbon of Ile360 contacts C7 of the phenyl ring and the apolar carbons of the Gln353 sidechain line the section of the pocket occupied by the relatively nonpolar fluoroethyl tail. Specific hydrogen bonding interactions also make prominent contributions to the binding of GTP-1. Lys353 lies at the base of the concave binding groove, where it forms a bifurcated hydrogen bond with the benzimidazole nitrogen (2.8 Å N–N distance) and the pyrimido nitrogen (3.4 Å) of GTP-1, satisfying the hydrogen bonding potential of the buried polar atoms within the tricyclic aromatic ring. Lys353 also completes its hydrogen bonding potential by forming a strong salt bridge with Asp358 in the same strand, and a weaker hydrogen bond with Asp358 in the adjacent strand. The oxygen of the Gln351 sidechain is well positioned to make a noncanonical hydrogen bond with the C–H bond of the beta carbon of the fluoroethyl tail, which points inward the fibril backbone. This tail orientation allows for close van der Waals contacts with backbone atoms in two strands and for the interaction with the sidechain of Gln351 (Fig. 2E). Overall, there is remarkable physiochemical and geometric complementarity between GTP-1 and the binding cleft of the tau filament.

Examining tau PHF:GTP-1, we observe that the GTP-1 heterocycles are situated at an optimal distance for pi-pi stacking (3.3–3.5 Å; fig. S10) (*33*), and GTP-1 forms an extended assembly scaffolded by the tau filament, reminiscent of supramolecular polymers which are highly cooperative (*34*). Unlike those molecules, GTP-1 contains both a rigid heteroaromatic and flexible nonaromatic region (aromatic: pyrimido[1,2-a]benzimidazole; nonaromatic: 2-fluoro-4-ethylpiperidine) (Fig. 3A). To assess the favorability of these interactions, we performed Hartree-Fock London Dispersion calculations (*35*). Each region of GTP-1 makes distinct contributions to the overall interaction; the major component (16 kcal/mol, 57%) indeed originates from the aromatic-aromatic interaction, whereas the smallest contribution comes from the cross interaction of the nonaromatic region with the aromatic region (5 kcal/mol, 18%), and the remainder comes from the nonaromatic-nonaromatic interaction (7 kcal/mol, 25%) (Fig 3B). Given that these subunits (aromatic and nonaromatic) have similar surface area (340 Å^2^ and 315 Å^2^, respectively), this speaks to the electronic favorability of stacking aromatic molecules as opposed to nonaromatic molecules. Moreover, this analysis does not consider entropic and hydrophobic contributions, which will also favor more rigid, aromatic molecules. The “tilt” angle of GTP-1, which leads to each compound crossing multiple tau strands, is congruent to that formed by the z-axis of the fibril, which is defined by a 4.77 Å repeat in this amyloid filament (note: helical twist is negligible over a short assembly) and the distance between two aromatic rings along the normal to the plane, typically most favorable between 3.3–3.5 Å. This angle is then described by a simple cosine relationship between these two distances, here 44° (Fig. 3, C and D). Given the commonality of these two constraints, we anticipate that adoption of a tilted heterocycle relative to the amyloid backbone may prove to be a common motif for binding filament structures, as this allows for significant favorable pi-pi interactions between small molecules while maintaining the translational symmetry of the amyloid.

**Fig. 3.**
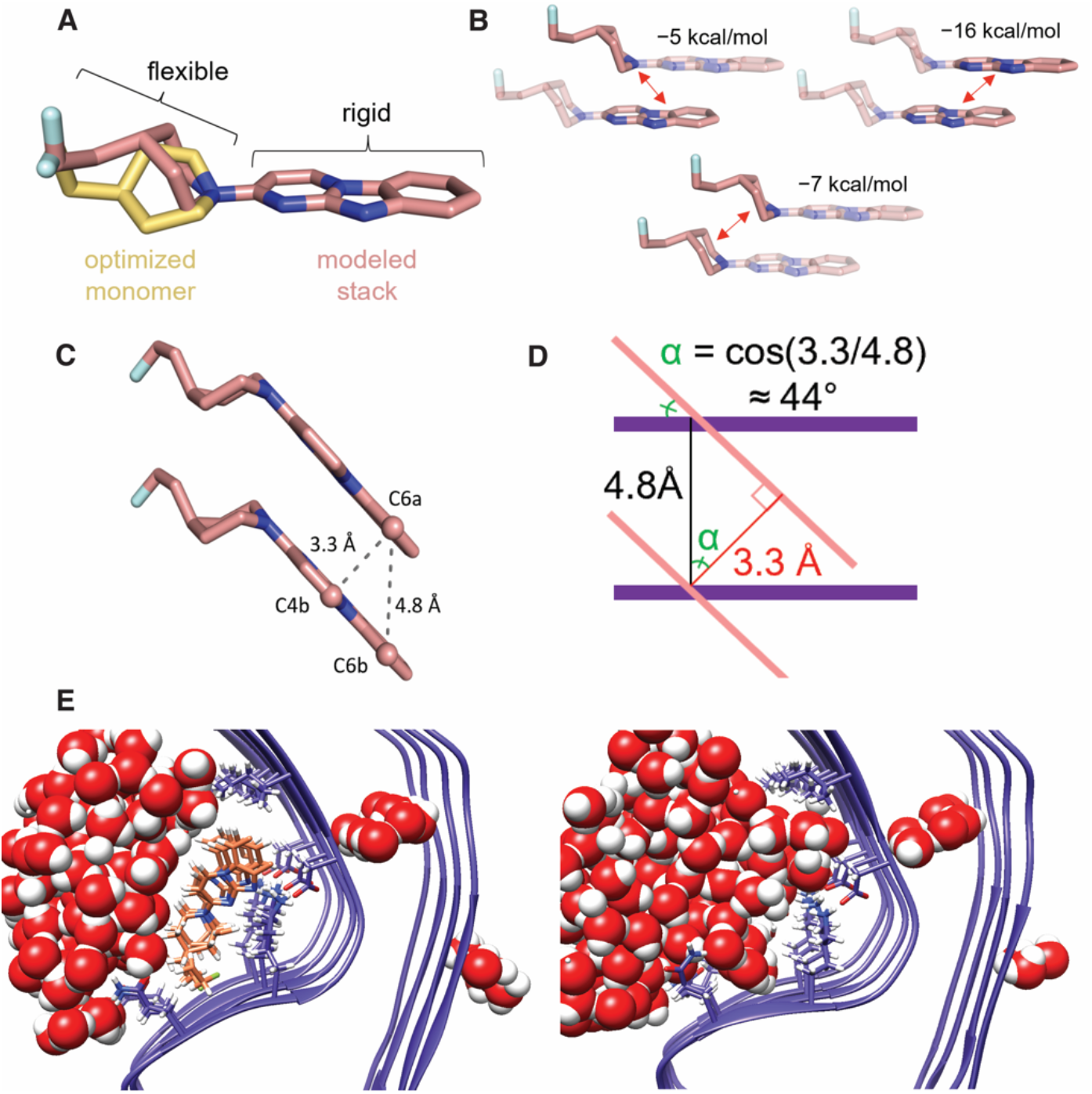
Favorable ligand-ligand interactions support stacked arrangement. (**A**) Comparison of the structure of GTP-1 monomer from an unconstrained DFT optimization (yellow) with the final modeled structure optimized in the context of amyloid-imposed constraints (coral). (**B**) Energy decomposition of the GTP-1 stacking interaction in a dimer using an HFLD calculation. (**C**) Illustration of the stacked GTP-1 interactions demonstrating the slipped nature of the stack, the retention of the amyloid displacement vector, and the distance of the pi-pi interactions. (**D**) Abstracted depiction of how the crossing angle between the plane of the amyloid backbone and the plane of the heterocycle is determined by the amyloid displacement vector and the optimal dimer interaction distance. (**E**) Representative final frames of a 100-ns MD simulation of tau PHF:GTP-1 (left) and unliganded tau PHF (right) demonstrate both the stability of the GTP-1 binding pose and the complete occlusion of water from the GTP-1 binding site throughout the trajectory.

The tau PHF:GTP-1 structure suggests a potentially powerful strategy for discovery and design of small molecules that bind with high affinity to amyloids in both a sequence- and conformation-specific manner. Filaments present a unique challenge for small-molecule design because their accessible surfaces tend to be relatively flat. This limits the amount of surface area potentially lost upon binding of a monomeric small molecule, hence the propensity for docking studies to show face-on binding of flat small molecules to the amyloid (*24, 25*). However, modeling of EGCG bound to one of the sites in tau PHFs by Eisenberg and colleagues indicates a similar ligand orientation to our tau PHF:GTP-1 model, in which the rings lie perpendicular to and match the symmetry of the fibril (*28*). Thus, this model, although at low resolution, suggests potential generality of this motif.

These structures suggest that this polymeric motif may have favorable filament binding properties, and we performed several calculations in an attempt to further examine this potential favorability. Although GTP-1 forms multiple productive contacts with the amyloid, the surface area lost upon binding a single monomer is negligible at 0.3 Å^2^. However, when two GTP-1 molecules stack, the overall loss of surface area increases to 85 Å^2^ (most of which is the apolar face of GTP-1) and creates a large driving force associated with the burial of hydrophobic groups as additional monomers are added. That this effect is not observed when two monomers are separated by an unliganded binding site suggests the system may be cooperative (table S3). To further examine this cooperativity, we undertook single-point density functional theory (DFT) calculations for binding of one, two, and three molecules of GTP-1 to five strands of a truncated model (residues 351–360) of tau (fig. S11). Although the accuracy of the calculations is intrinsically limited due to their static nature and lack of explicit solvation, potential trends can be gleaned. Notably, the binding energy of a single tracer against the five strands is the same in all three potential binding sites. For two tracers bound in adjacent sites, the energy is the sum of the small molecule–protein binding energies and the small molecule–small molecule dimerization energy, suggesting positive cooperativity. The same trends continue with three tracers (the minimal model for an extended stack), suggesting the calculations are relevant to the overall assembly. In contrast, two tracers separated by an unliganded binding site (a minimal model for sparse binding) shows no favorable small molecule–small molecule binding energy (table S3). We then used molecular dynamics to simulate five stacked ligands centered in nine strands of both protofilaments and found both the tau filament and the stacked assembly to be stable over 100 ns (fig. S12). Throughout the simulation, the GTP-1:tau and GTP-1:GTP-1 interactions seen in the experimental structure were maintained, and no penetration of water was observed into the dry protein–small molecule interface, confirming the geometric and electrostatic complementarity of stacked GTP-1 with this binding groove (Fig. 3E).

Moreover, the observed behavior, that both small molecule–protein and small molecule–small molecule interactions are local and that the latter are positively cooperative, is analogous to other well-studied biological systems. These systems, including the random coil to helix transition of a polypeptide or the binding of dye molecules to DNA, are well described by mathematical models (*36–38*). This suggests a route forward to better understanding the thermodynamic and kinetic behavior of small molecule–amyloid interaction under physiological conditions. In addition to the tau PHF:EGCG structure (*28*), templated assembly and symmetry matching have also been observed in the assemblies of similar aromatic molecules with globular proteins, although the limited size of the binding sites restricts the assembly size to a maximum of four molecules (*39–41*).

Rather than binding to a nondescript surface along a uniform β-sheet, the strong geometric and physical complementarity between GTP-1 and this unique cleft likely imparts considerable specificity for AD filaments (Fig. 4). The local architecture of Gln351 to Ile360 that comprises the GTP-1 binding site is markedly different in filament structures of other tauopathies. In many cases, the key residues that form close contacts in the AD structure are either not solvent exposed or instead form a convex surface as opposed to the concave cleft suitable for binding. Although chronic traumatic encephalopathy (CTE) protofilaments have a similar C-shaped architecture to AD, this region of the CTE filament structure is defined by a much shallower angle formed by the kink at Gly355. This causes Ile360 to shift ~3 Å further from Gln351 than in the AD structure, resulting in the loss of the apolar interaction between Ile360 and C7 of the GTP-1 phenyl ring (fig. S13). Based on these structural differences, GTP-1 likely does not stack in this cleft of CTE filaments. While it is possible that GTP-1 binds to other β-sheet folds, it would likely involve an alternate mode of binding and different sequence elements within the tau filament structure.

**Fig. 4.**
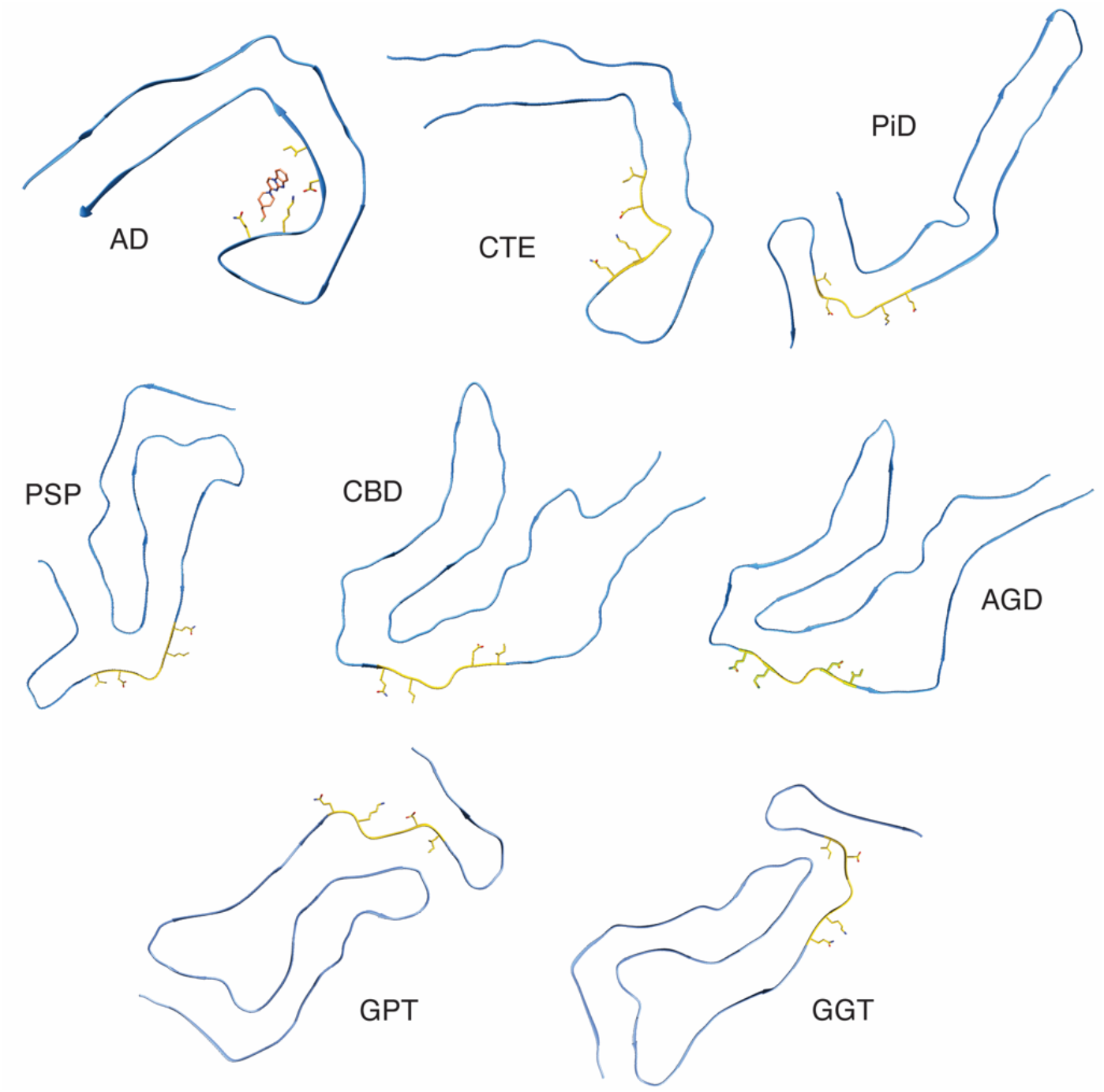
Comparison of the GTP-1 binding pocket residues in other tau filament structures. The ligand binding pocket of GTP-1 is highlighted in gold, and the specific residues forming the binding pocket (Gln351, Lys 353, Asp358, and Ile360) are shown. This binding pocket is unique to AD filaments compared to existing filament structures, thus indicating GTP-1 binding may be specific to the AD conformation.

Symmetry matching, as observed in the structure of GTP-1 bound to PHFs from a patient with AD, may provide a powerful strategy to increase the druggability of available binding sites in filaments. In an emergent system such as this, small changes to the binding site likely confer a large effect on the binding of GTP-1. Thus, designing small-molecule compounds with high specificity and affinity for a single site within the amyloid filament conformation may be feasible. This analysis suggests that in the development of future tools for diagnostics and, potentially, therapeutics, an emphasis should be placed on heterocycles that stack favorably in the context of the amyloid axial symmetry and on achieving shape and electrostatic synergy with the targeted binding cleft. Understanding not only the amyloid assembly as a supramolecular entity, but also the small molecule, reveals a new route to designing amyloid filament binding small molecules.

## Acknowledgements

We thank Bill Seeley and the UCSF Neurodegenerative Disease Brain Bank for providing patient tissue for this study. We also thank Neil Vasdev and MedChem Imaging for providing GTP-1 PET ligand.

## Funding

This work was funded by the Rainwater Charitable Foundation (D.R.S.), the National Institutes of Health (P01AG002132: D.R.S., S.B.P., and W.F.D.; F32GM139379: M.J.C.). The UCSF Neurodegenerative Disease Brain Bank receives funding support from NIH grants P30AG062422, P01AG019724, U01AG057195, and U19AG063911, as well as the Rainwater Charitable Foundation and the Bluefield Project to Cure FTD.

## Author contributions

G.E.M. purified patient tissue samples, prepared EM grids, collected data, performed cryo-EM image analysis, performed model building and refinement, developed figures, and wrote and edited the manuscript. M.J.C. performed calculations, performed model building and refinement, developed figures, and wrote and edited the manuscript. S.T. performed calculations and developed figures. E.T. operated the Krios microscope and helped with data collection. J.L. performed tau quantification and infectivity assays. S.B.P. supervised the project and edited the manuscript. N.A.P. provided critical ideas and methods and edited the manuscript. W.F.D. supervised the project and edited the manuscript. D.R.S. initiated the project, supervised the project, and edited the manuscript.

## Competing Interests

S.B.P. is the founder of Prio-Pharma, which did not contribute support for this study. W.F.D. is a member of the scientific advisory boards of Alzheon Inc., Pliant, Longevity, CyteGen, Amai, and ADRx Inc., none of which contributed support for this study.

## Data and Materials Availability

Cryo-EM maps and atomic coordinates have been deposited in the EMDB and PDB with accession codes: EMD-XXXX and PDB YYY.

