## Supplementary Materials for "Stacked binding of a small molecule PET tracer to Alzheimer’s tau paired helical filaments"

**This PDF file includes:**

Materials and Methods
Supplementary Text
Figs. S1 to S13
Tables S1 to S3

### Materials and Methods

Purification of tau filaments

Filament purification was based on Fitzpatrick *et al.*, 2017 (*10*). Briefly, 5 g of fresh-frozen frontal cortex tissue from an 88-year-old male patient with Alzheimer's disease was homogenized at 10 mL/g of tissue in 10 mM Tris-HCl (pH 7.4), 800 mM NaCl, 1 mM EGTA, and 10% sucrose. The homogenate was centrifuged at 20,000 g for 10 minutes, and the supernatant was kept. The pellets were resuspended in 5 volumes of the same buffer, centrifuged again, and the 2 supernatants were combined. A final concentration of 1% N-laurosarcosinate (w/v) was added to the combined supernatant, and this mixture was incubated for 1 h at room temperature. It was then centrifuged at 100,000 g for 1 hour, and the pellets were resuspended in 30 volumes of 10 mM Tris-HCl (pH 7.4), 800 mM NaCl, 1 mM EGTA, 5 mM EDTA, and 10% sucrose. This was followed by another centrifugation at 20,100 g for 30 minutes at 4°C. The supernatant was kept and centrifuged at 100,000 g for 1 hour, and the final pellet resuspended in 20 mM Tris-HCl (pH 7.4) and 100 mM NaCl at a concentration of 10 μL/g frozen tissue.

Negative stain imaging

Purified frontal cortex tissue was diluted 1:10 for a final concentration of 100 μL/g frozen tissue. 5 μL was added to a glow-discharged 400 mesh copper grid with a layer of amorphous carbon. After 30 seconds, the grid was blotted with filter paper, washed, and blotted twice with nanopore water. 5 μL of 0.75% uranyl formate was then added and blotted. Three more 5 μL aliquots of uranyl formate were added and removed by vacuum aspiration. Images were collected on a Talos L120C (Thermo Fisher Scientific) operating at 200 kV and equipped with a Ceta-D (Thermo Fisher Scientific) camera.

Tau quantification and infectivity

Total tau in the final purification fraction frozen on grids was quantified using a Total Tau cellular kit (HTRF, Cisbio). The output was read on a PHERAstar FSX plate reader (BMG LABTECH). A standard curve was generated using recombinant 0N4R tau.

Infectivity assays were then performed similarly to Woerman *et al.*, 2016 (*30*), and an HEK293T cell line expressing the 4R repeat domain of tau (residues 243 to 375 in 2N4R tau) with mutations P301L and V337M fused to yellow fluorescent protein (YFP) at the C-terminus was used. Cells were cultured and plated in 1x Dulbecco's modified Eagle medium supplemented with 10% (vol/vol) fetal bovine serum. Cells were plated in a 96 well plate (3,000 cells per well) with 0.1 μg/mL final concentration of Hoechst 33342. Cells were then returned to an incubator that maintained a humidified atmosphere of 5% $CO_2$ at 37°C for 2 hours. Samples were diluted to the appropriate tau concentration with Dulbecco's PBS (DPBS), mixed with Lipofectamine 2000 (Thermo Fisher Scientific; final concentration: 0.03%), and incubated for 1 hour at room temperature. Samples were then added to the cells in six replicate wells and incubated for 3 days at 37°C in a humidified atmosphere of 5% $CO_2$. Plates were imaged using the IN Cell Analyzer 6000 cell-imaging system (GE Healthcare). Images were then analyzed using the IN Cell Developer software (GE Healthcare), with an algorithm that detects aggregated protein using pixel intensity and size thresholds in living cells. The output, DxA, is a measure of the size and brightness of these aggregates.

Cryo-EM grid preparation and data collection

Purified frontal cortex tissue was incubated with 20 μM ligand for 45 minutes prior to freezing. Three μL of this mixture was added to a glow-discharged 200 mesh 1.2/1.3R Au Quantifoil grid for 10 seconds before blotting for 2 seconds. A second 3 μL aliquot was added for 3 seconds and blotted for 1 second before being plunge frozen in liquid ethane using a FEI Vitrobot Mark IV (Thermo Fisher Scientific). Super-resolution movies were collected at a nominal magnification of 105,000x (physical pixel size: 0.417 Å per pixel) on a Titan Krios (Thermo Fisher Scientific) operated at 300 kV and equipped with a K3 direct electron detector and BioQuantum energy filter (Gatan, Inc.) set to a slit width of 20 eV. A defocus range of 0.8 to 1.8 μm was used with a total exposure time of 2.024 seconds fractionated into 0.025 seconds subframes. Movies were motion-corrected using MotionCor2 (*42*) and were Fourier cropped by a factor of 2 to a final pixel size of 0.834 Å per pixel.

Image processing

For GTP-1, 15,160 micrographs were collected, and all processing was done in RELION 3.1 (*43*). Dose-weighted summed micrographs were imported into RELION 3.1. The contrast transfer function was estimated using CTFFIND-4.1. Filaments were manually picked and then segments were extracted with a box size of 900 pixels downscaled to 300 pixels. A larger box size of 1,200 pixels downscaled to 300 pixels was used to estimate the filament crossover distance. Contaminants and segments contributing to straight filaments were separated out using reference-free 2D class averaging. The remaining segments were re-extracted with a box size of 288 pixels without downscaling. The map from EMDB 0259 (*12*) low-pass filtered to 15 Å was used as an initial model. One or more rounds of 3D classification with image alignment were performed, with helical rise and tilt parameters fixed to eliminate obvious junk particles. Local rise and tilt were fixed during a first round of 3D auto-refinement using $C_1$ symmetry and a PHF map low-pass filtered to 10 Å. A second round of 3D auto-refinement was run imposing $C_{2_1}$ symmetry and allowing rise and twist parameters to vary, using the map from the first auto-refinement low-pass filtered to 4.5 Å as a model. Contrast transfer function (CTF) refinement was then run, fitting the defocus and astigmatism, as well as estimating 4$^{th}$ order aberrations. These particles were then used in a 3D classification job allowing the rise and twist to vary, but without image alignment. Particles contributing to the highest resolution map(s) were selected, and a final 3D auto-refinement was run. Maps were sharpened using the standard post-processing procedures in RELION. Full statistics are shown in table 1.

A reconstruction of straight filaments was attempted using the same workflow, but a high-resolution structure was unable to be obtained, even after extensive 2D and 3D classification.

Refinement of Tau PHF

Prior to ligand placement, a single strand of a previously solved PHF model (PDB: 6HRE) (*12*) was refined against the density using Phenix (*44*). Refinement of side chains in the GTP-1 binding pocket was done in COOT (*45*). This apo model was then translated to give a stack spanning 5 rungs and validated in Phenix.

Computation and modeling of GTP-1:

*General Considerations:*

All molecular mechanics-based conformer searches were performed using the ConfGen tool in Maestro (*46*). The OPLS4 forcefield (*47*) was used, and an energy threshold of 21 kJ/mol (or 5.02 kcal/mol) was used. All density functional theory (DFT) calculations were performed using ORCA 5.0.3 (*48*). Optimizations were performed using the BP86 functional (*49, 50*) and the def2-SVP basis sets (*51*) with an auxiliary basis set approximation (*52*), a dispersion correction (*53*), and a solvent polarization model (CPCM) (*54*). Dichloromethane was used as the solvent model because it has dielectric properties similar to those found in proteins (*55*). In cases of unconstrained optimization, a numerical frequency calculation was performed to confirm that the geometry was at a global minimum. In cases where a constrained optimization was performed, the electronic energy was used for comparison as an estimate of the enthalpy, which is valid assuming a similar zero-point energy, vibrational energy, rotational energy, and translational energy. This estimate is necessary because these systems are not at a global minimum, and, thus, the exact calculation of the enthalpy and entropy via DFT will be prone to errors. Given that systems of similar size are being compared, the following estimate should be valid for determining relative energies:

$$U = E_{\text{electronic}} + E_{\text{zero point energy}} + E_{vibrational} + \mathrm{E}_{rotational} + \mathrm{E}_{translational} \quad \text{(Eq S1)}$$

$$H = U + \mathrm{k_B T} \quad \text{(Eq S2)}$$

*Modeling of GTP-1:*

A DFT-minimized monomer of GTP-1 was used as the input for the initial conformer search (0.5 Å RMSD) in Maestro. Outputs (43) were clustered by the position of the piperidine ring (maximum atom distance <0.5 Å) ignoring the fluoroethyl tail. The centroids and their fit to the density can be seen in fig. S9. From the best fit conformer, a dimer was then generated taking into account the translational vector of the amyloid (the rotational element is considered to be negligible over two units). The dimer was then optimized for a series of torsional angles and the electronic energies were compared (fig. S10). The lowest energy torsion also improved the fit to the density, so that was used for a further conformational search in Maestro. In this search, all of the atoms of the tricyclic aromatic and the piperidine ring were held constant (i.e., only the fluoroethyl tail was varied) and the outputs were required to have at least one atom that varied by more than 0.1 Å. Both small molecule–small molecule and small molecule–protein clashes were then considered for the outputs (13). Clashes were defined as two heavy atoms (C, N, O, F) with a distance of <2.5 Å. All outputs that passed the clash filter (5) were again subjected to a constrained DFT optimization, and that final output was compared to the cryo-EM density. Selecting the best output based on the density, a final refinement was done in Phenix.

For the modeled conformer, the interaction energy can be evaluated via a Hartree–Fock London Dispersion calculation, which decomposes the overall energy of a system into the energy of the individual units and the energy arising from their interactions. This calculation was performed using the def2-TZVP(-f) basis set (*51*) with the auxiliary basis sets def2/J (*52*) and def2-TZVP/C (*56*) in a continuous polarized solvent model (*52*). The interaction energy of −26 kcal/mol for a GTP-1 dimer can be decomposed into the components coming from the aromatic and nonaromatic subregions by performing calculations on those individual pairs with a proton capping the portion of the molecule that was removed.

*Binding to the Amyloid:*

Estimates of the interaction energy of the small molecule(s) with the protein could be readily achieved via single point calculations in the apo- and holo-state. To speed calculations, a truncated active site region was considered, consisting of residues 351–360 of a given strand with protons added to cap the backbone. Five strands in total were considered. A single PET tracer appears to interact with three strands of the amyloid backbone via visual inspection, so there were three binding sites across the five strands. Single point calculations were performed with a single GTP-1 in each of the binding sites, and the energies were confirmed to be constant, suggesting that protein–small molecule interactions are local (table S2). If long-range interactions were observed, then positioning GTP-1 in the middle binding site should be more favorable. This validates the model size.

We also performed calculations in which GTP-1 occupies both the top two or both the bottom two binding sites, which were isoenergetic. However, calculations with two GTP-1 spaced out (i.e., the middle binding site is empty) show a lower energy. A final calculation in which all three sites are occupied by GTP-1 confirms the trends shown with two sites. As every additional GTP-1 added after three effectively introduces another unit into the interior of the stack, a calculation on a larger system is not needed.

The $\Delta\Delta E_{\text{binding}}$ term (table S2) is evaluated by comparing the energy to the energy of binding one GTP-1 in the middle of the stack adjusted for the stoichiometry. The negative terms seen for the 2 GTP-1 (top), 2 GTP-1 (bottom), and 3-GTP-1 are a result of the cooperative effect of the GTP-1 interactions. The magnitude of that interaction (about −19 kcal/mol) suggests that DFT probably slightly underestimates this dispersion-based interaction, a well-known phenomenon (*57*).

The surface area was evaluated using the same truncated systems with the get area feature in PyMOL (solvent turned on and the dot density set to the maximum) (*58*).

### Atomistic Molecular Dynamics Simulations of Ligand-bound Tau

*Simulation Parameters and Analysis:*

The MD system was prepared using AmberTools in Amber18 (*59, 60*). The N-termini of the cryo-EM structure were acetylated, while the C-termini were amidated. The electrostatic potential of GTP-1 was calculated using Gaussian09 (*61*), which was subsequently used to fit partial charges of the molecule. Additional GTP-1 parameters were generated using AmberTools. The structure was then solvated with SPC/E-modeled waters in an octahedron with 8Å buffer from the protein, and the system was neutralized by adding Cl- ions. All simulations were performed using Amber18 with the ff14SB forcefield (*62, 63*). Simulations began with 1,000 restrained steepest-descent minimization steps before switching to a maximum of 5,000 steps in conjugate gradient steps. The system was then heated up to 300K over 50 ps in NVT equilibration with Langevin thermostat control of temperature and harmonic restraints on protein and small molecule atoms with a 10 kcal/(mol·$Å^2$) force constant. The system was then switched to NPT, which used the Monte Carlo barostat to maintain pressure at 1 atm. The restraints were gradually removed over 1 ns, and the simulation progressed to an unrestrained production run for 100 ns.

The systems were simulated under periodic boundary conditions, employing the SHAKE algorithm with 2.3 fs timesteps. Particle Mesh Ewald was used for long-range electrostatics, and non-bonded interactions were cut off at 8Å. Two independent simulations of GTP-1–bound paired helical filaments were performed, for a total of four simulated protofilaments. Using MDAnalysis (*64, 65*), time series of RMSDs were calculated to the starting cryo-EM structure as a measure of conformational stability.

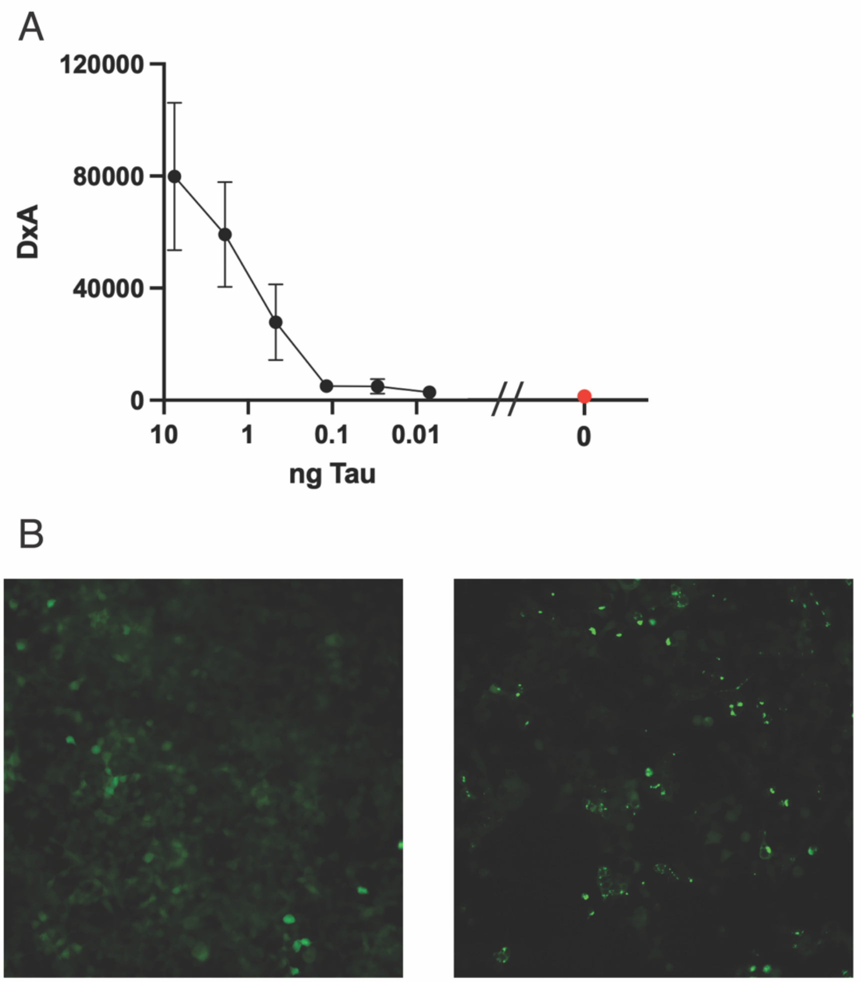


**Fig. S1. Infectivity of tau filaments purified from human AD brain tissue for cryo-EM structure determination.**
Infectivity of the partially purified tissue from which the GTP-1 co-structure was solved, as measured by a cell-based fluorescence assay. The tissue was incubated for 3 days with HEK-293T cells that express the repeat domain of 4R tau–containing mutations P301L and V337M fused to YFP (*66*). The level of infectivity was measured by the size and brightness of fluorescent puncta formed in the cells, which is quantified as DxA (see Methods). (**A**) Quantification of partially purified tissue infectivity over a range of total tau in the sample, as determined using a total tau homogenous time-resolved fluorescence (HTRF) assay (*67*). As increasing amounts of tau were added to the cells, the DxA increased, indicating that our imaged sample was pathogenic and disease relevant. The red point represents a control with no tau added. (**B**) Representative images from a control with no tau (left) and a sample incubated with 7.5 ng total tau (right). The diffuse fluorescence in the left image indicates a lack of infectivity, while the distinct puncta in the right image are characteristic of pathogenicity.

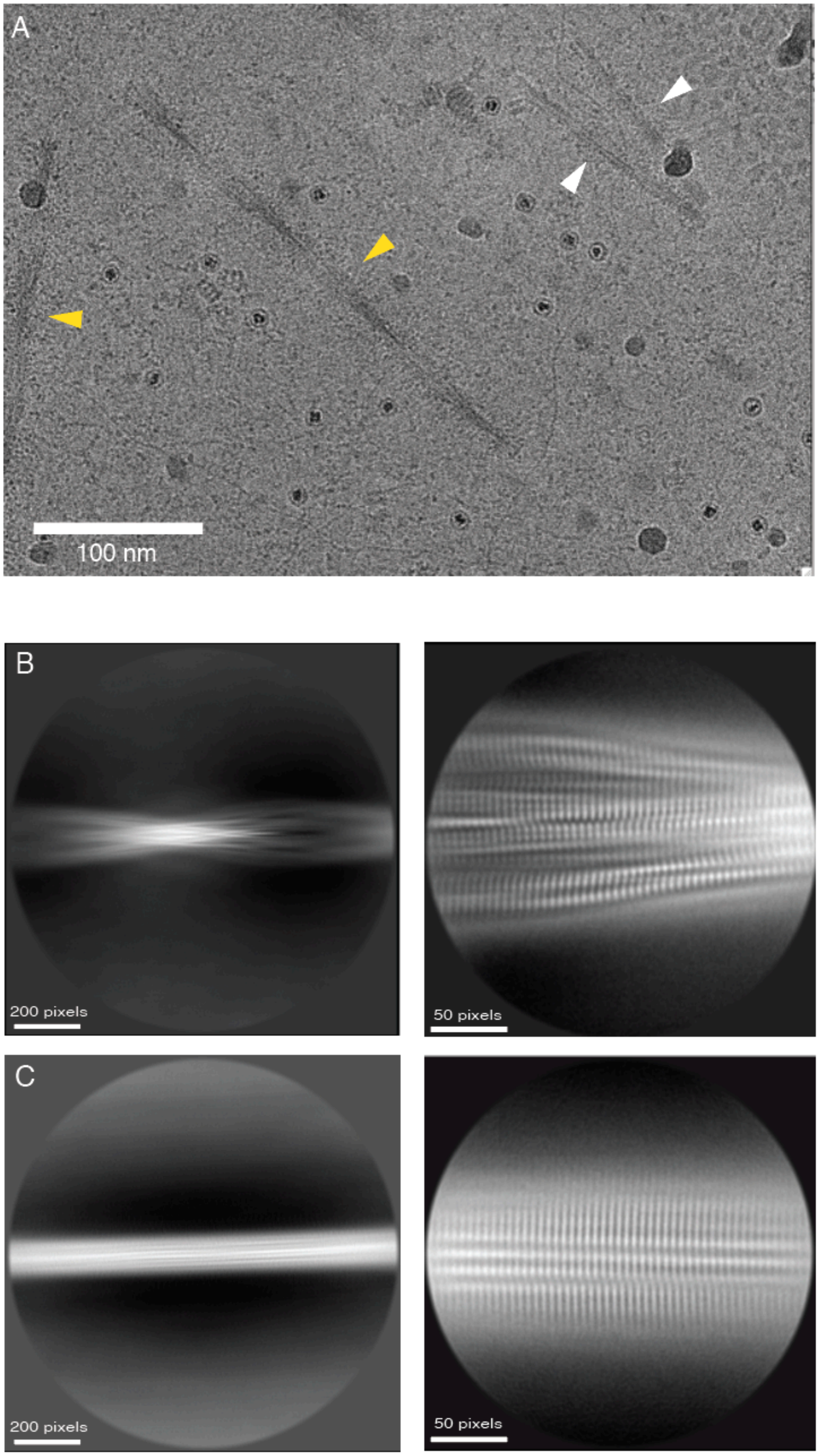

A
100 nm
B
200 pixels
50 pixels
C
200 pixels
50 pixels

**Fig. S2. Micrograph and 2D class averages of AD filaments used for 3D structure determination.**

(**A**) Representative cryo–electron micrograph showing both paired helical filaments (PHFs) and straight filaments (SFs) purified from AD patient tissue. PHF (gold arrows) and SF (white arrows) determination is based on crossover distance and comparisons to previous image data (*10*). Representative reference-free 2D class averages are shown for (**B**) PHFs and (**C**) SFs, with box sizes 1,200 pixels downscaled to 300 pixels (left) and 288 pixels without downscaling (right).

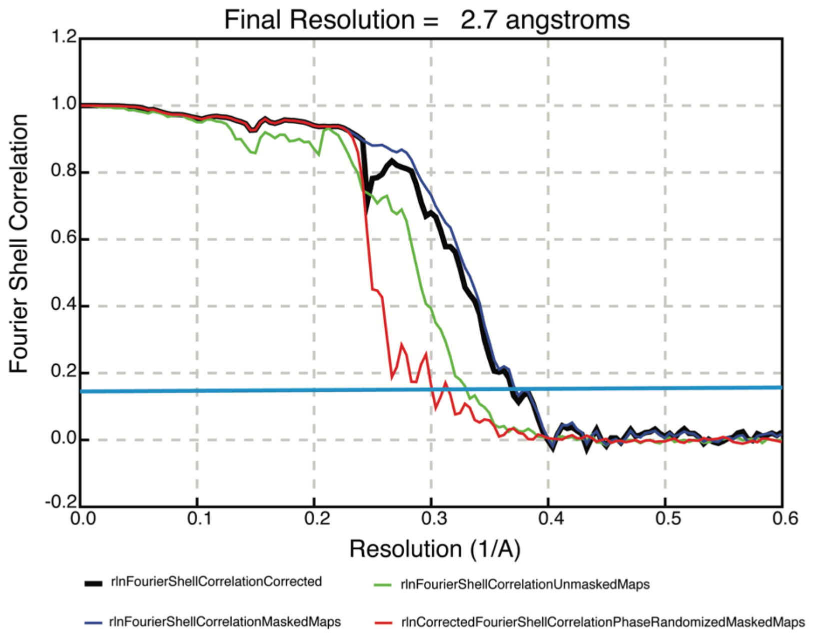


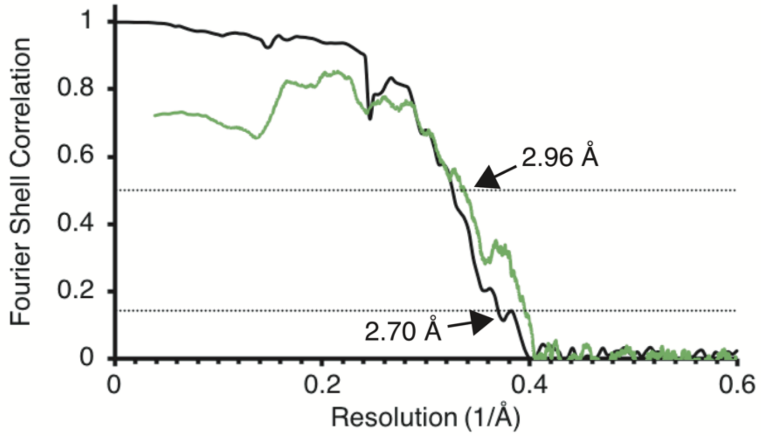


**Fig. S3. Fourier shell correlation (FSC) curves.**
(**A**) FSC curves for two independently refined cryo-EM half maps of the tau PHF:GTP-1 structure. (**B**) Corrected FSC curve from (A) in black and the FSC curve for the refined atomic model against the final cryo-EM map in green.

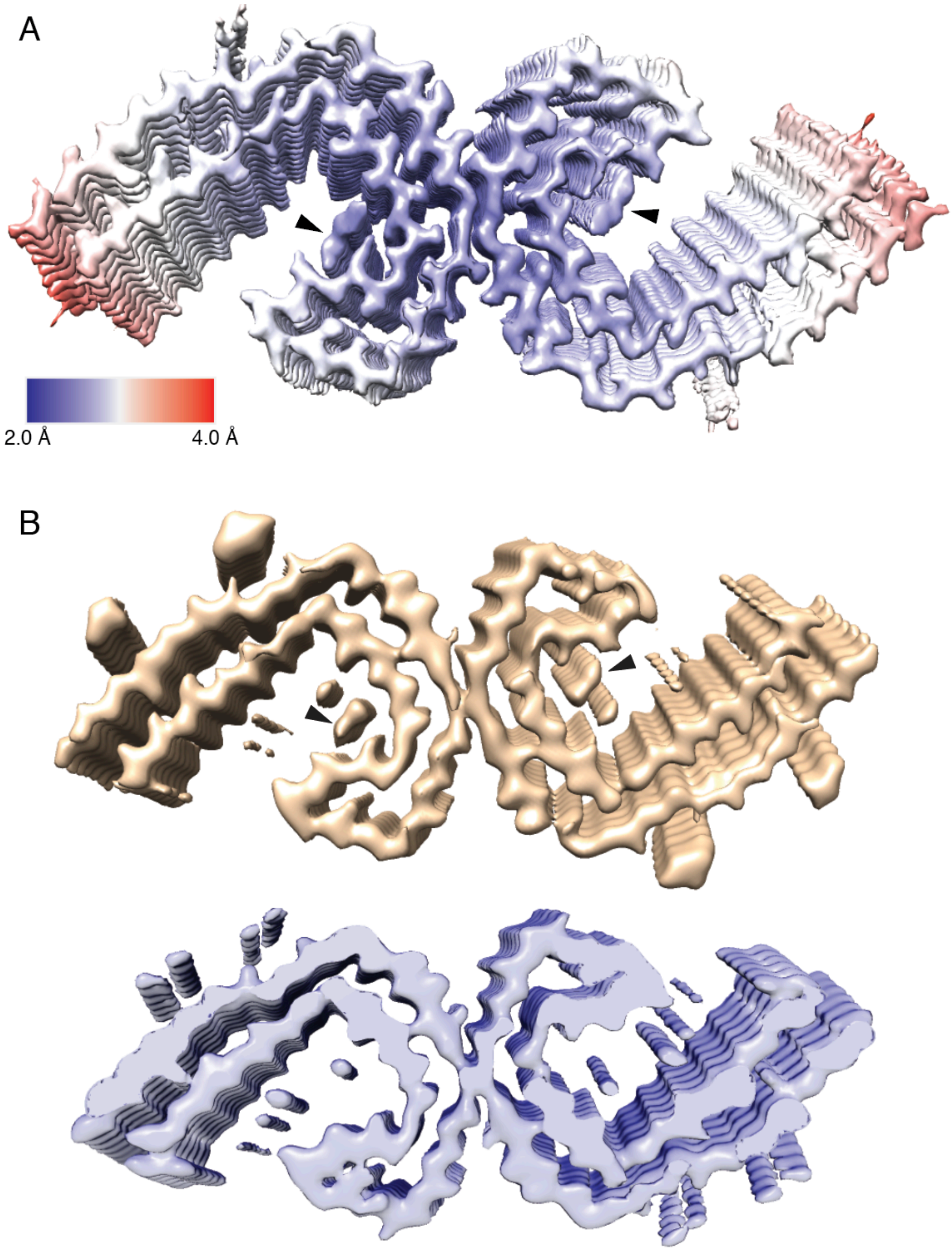
A
2.0 Å
4.0 Å
B

**Fig. S4. Local resolution estimation and comparison of PHF:GTP-1 with apo tau PHF (EMD-0259) cryo-EM maps.**

(**A**) Local resolution map of tau PHF:GTP-1 showing high resolution (~2.5 Å) of the filament core and GTP-1 ligand (arrow). (**B**) Cryo-EM maps of tau PHF:GTP-1 (top: gold), in comparison to the previously solved structure (EMD-0259; bottom: blue) low-pass filtered to 5 Å. Additional density ascribed to GTP-1 is indicated (arrow). No densities unique to the PHF:GTP-1 map are identified, indicating specific binding.

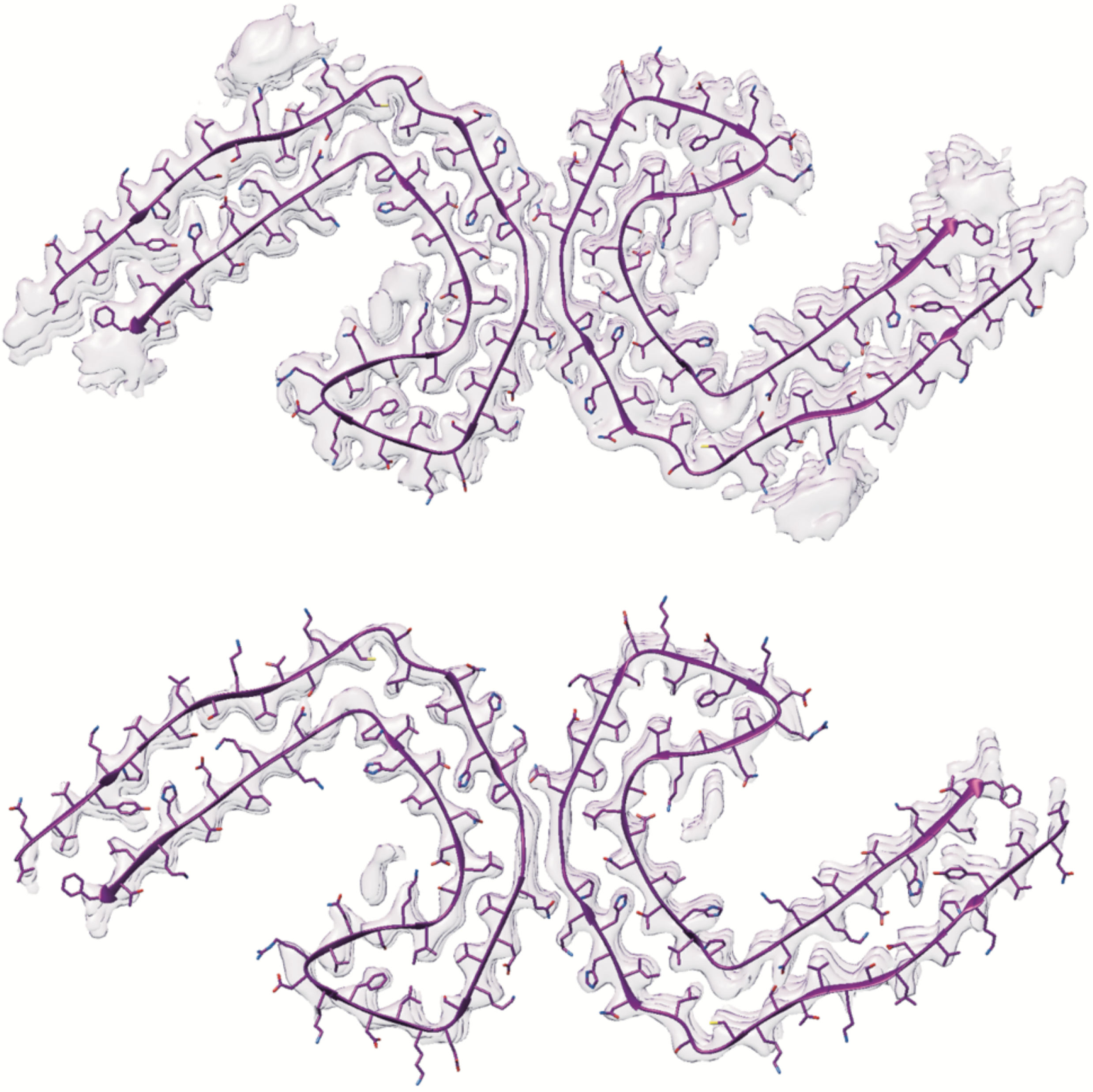

**Fig. S5. PHF + GTP-1 cryo-EM map at two thresholds.**
Cryo-EM map of AD PHFs in complex with GTP-1 low-pass filtered to 3.5 Å at Chimera threshold level 0.0095 (top, σ = 3.0) and 0.0243 (bottom). While other densities surrounding the amyloid filament disappear at high threshold, the density corresponding to GTP-1 remains, indicating high binding occupancy.

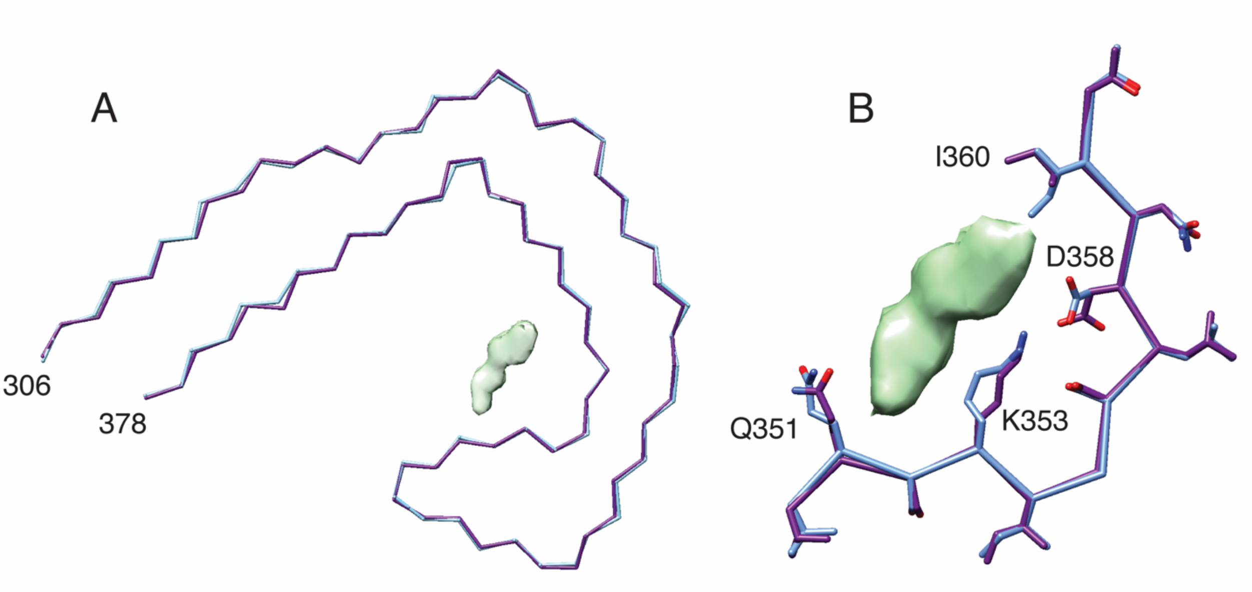


**Fig. S6. Comparison of GTP-1 bound and apo tau PHFs.**
(**A**) Overlay of the backbone structures of PHFs with (purple) and without (blue) GTP-1 bound.
(**B**) Overlay of the residues in the GTP-1 binding pocket with (purple) and without (blue) GTP-1 bound, showing subtle sidechain rearrangements.

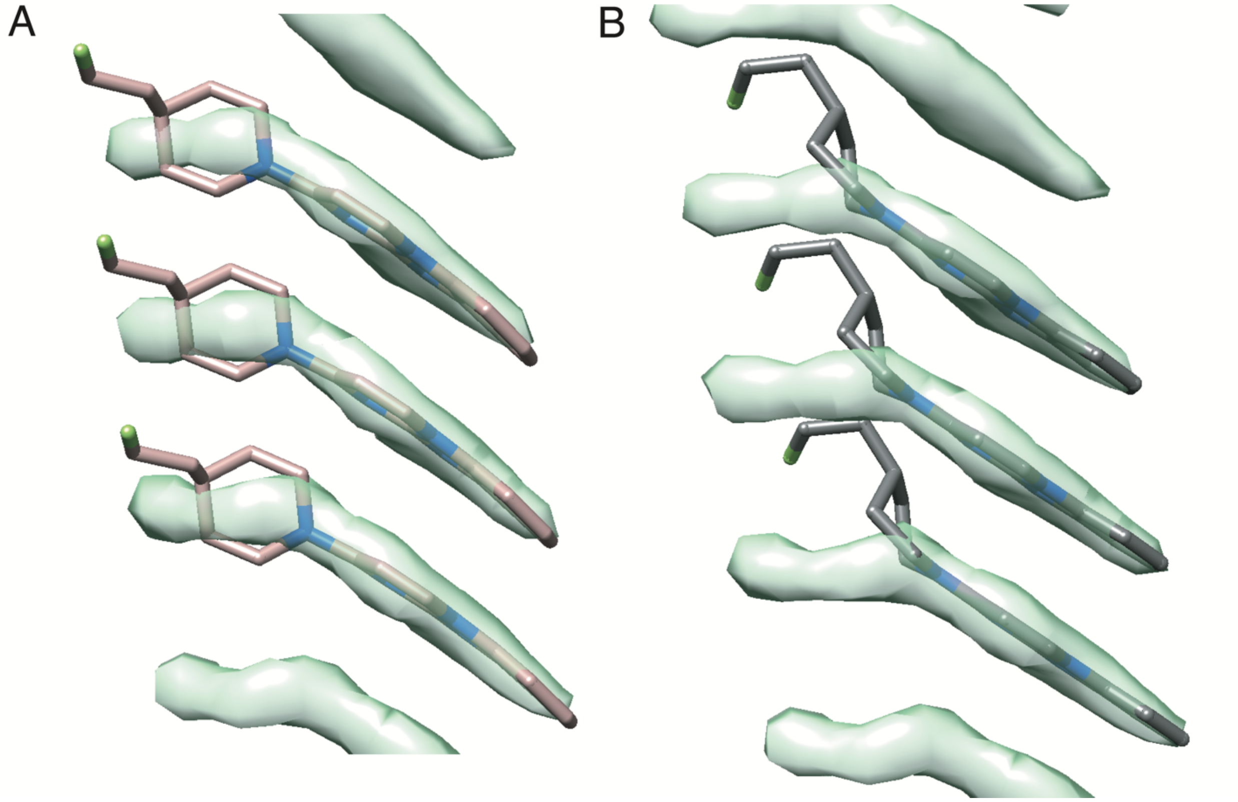


**Fig. S7. Map+model of GTP-1 comparing Phenix and DFT optimization modeling approaches.**

GTP-1 monomer conformations generated by (**A**) PHENIX eLBOW and (**B**) DFT optimization of a monomer fit into the ligand density. The distance of closest approach between stacked molecules is 2.3 Å for the eLBOW conformation and 1.7 Å for the DFT conformation.

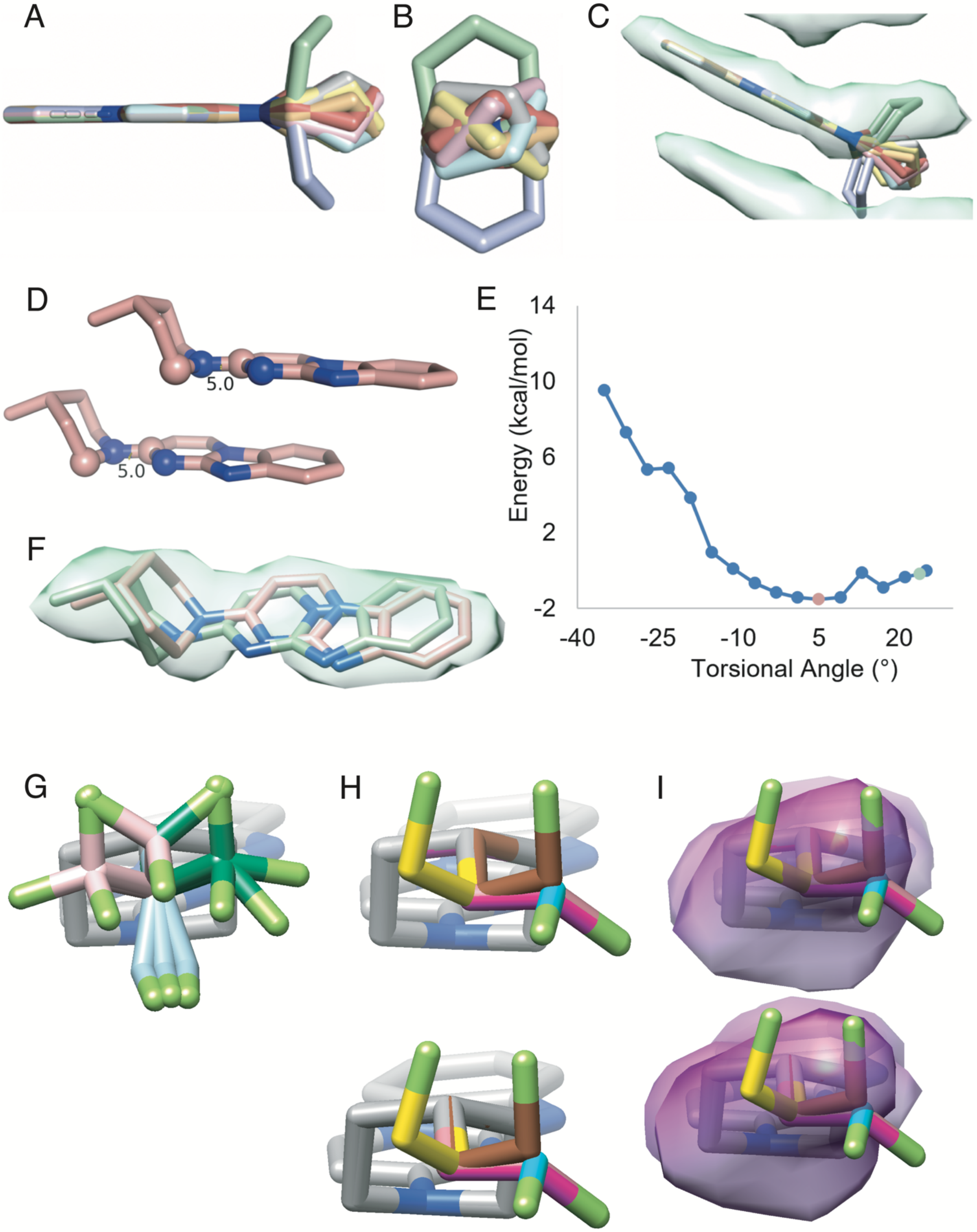
A
B
C
D
5.0
5.0
E
14
10
6
2
-2
-40
-25
-10
5
20
Energy (kcal/mol)
Torsional Angle (°)
F
G
H
I

**Fig. S8. Conformational search of the flexible nonaromatic region of GTP-1.**
(**A**–**C**) Conformational search of the piperidine ring. The possible conformers are related by one mirror plane in the plane of the tricycle and a pseudo-mirror plane that is orthogonal to the plane of the tricycle and lies along the C–N bond connecting the tricycle to the piperidine ring. (**A**) Edge-on view of centroids resulting from piperidine ring search. Centroids generated with a maximum atom distance of 0.5 Å. (**B**) End-on view of piperidine ring centroids. (**C**) Comparison of the centroids placed into the density, which clarifies that the light green centroid is the correct ring conformation. (**D**–**F**) Torsional search of the piperidine ring. The potential energy surface is very soft except for where clash occurs with the dimer. The energy minimum presumably arises from better donation of the piperidine nitrogen into the aromatic tricycle. (**D**) Example of one output from the constrained DFT optimizations. The torsional angle defined by the atoms indicated as spheres were constrained. Also, the translational distance between all of the atoms in the piperidine ring was constrained across the dimer. (**E**) A graph of the energy landscape for the torsional angle. The green dot was the Maestro output (23.8°), and the coral dot (5°) was the minimum energy angle found. (**F**) The starting Maestro conformation (green) and the torsion-optimized conformation (coral) in the cryo-EM density showing the improved fit upon torsional optimization. (**G**–**I**) Conformational search of the fluoroethyl tail. In the absence of the protein and the small molecule stack, the fluoroethyl conformers occupy essentially a three-fold symmetric well with a pseudo-mirror plane that is along the C–N bond connecting the tricycle to the piperidine ring and that is orthogonal to the tricycle. (**G**) All of the outputs from the search in Maestro using 0.1 Å atom maximum distance. The conformers in light blue clashed with their dimeric partner, the conformers in pink clashed with the protein, and the conformers in green were taken forward for constrained optimization as dimers. (**H**) The dimers that resulted from constrained optimization of the Maestro outputs as dimers using a translational constraint on the piperidine ring and fluoroethyl tail atoms. (**I**) Dimer outputs compared to the density. The bright brown monomer was used as the input for the final Phenix refinement.

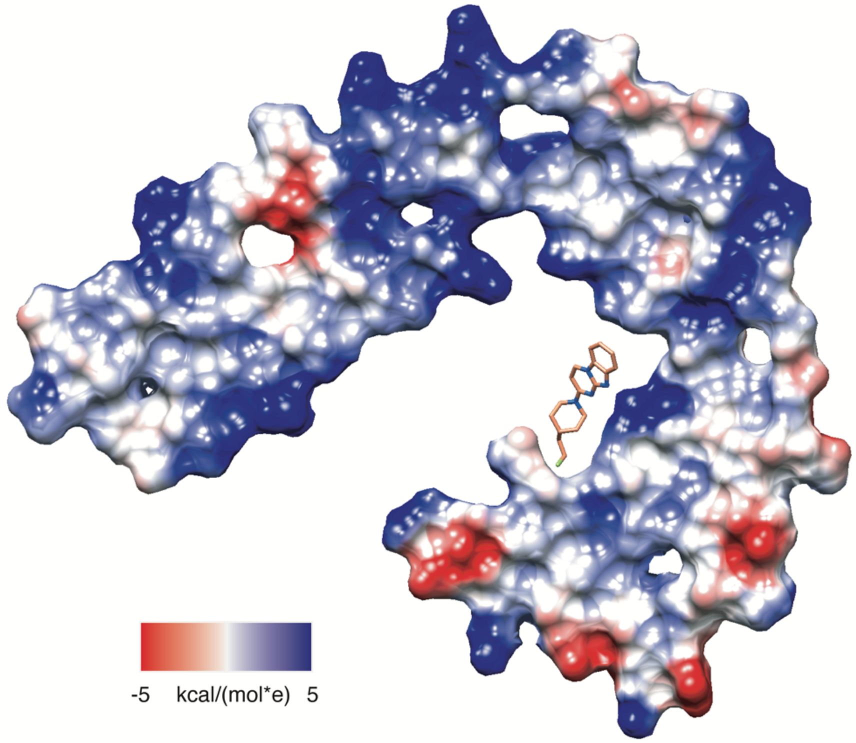


**Fig. S9. Solvent accessible surface of an AD PHF with GTP-1 modeled.**
GTP-1 binds in a cleft in the AD PHF that shows strong geometric and electrostatic complementarity to the small molecule.

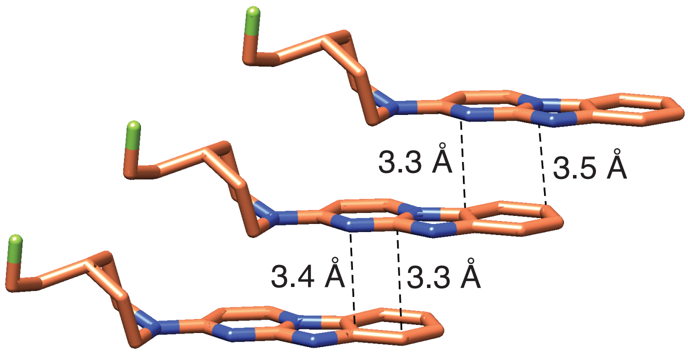


**Fig. S10. Distance between the heterocycles in a GTP-1 stack.**
Distance measurements between stacked GTP-1 molecules showing that the heterocyclic portion of GTP-1 is situated at an optimal distance for pi-pi interactions.

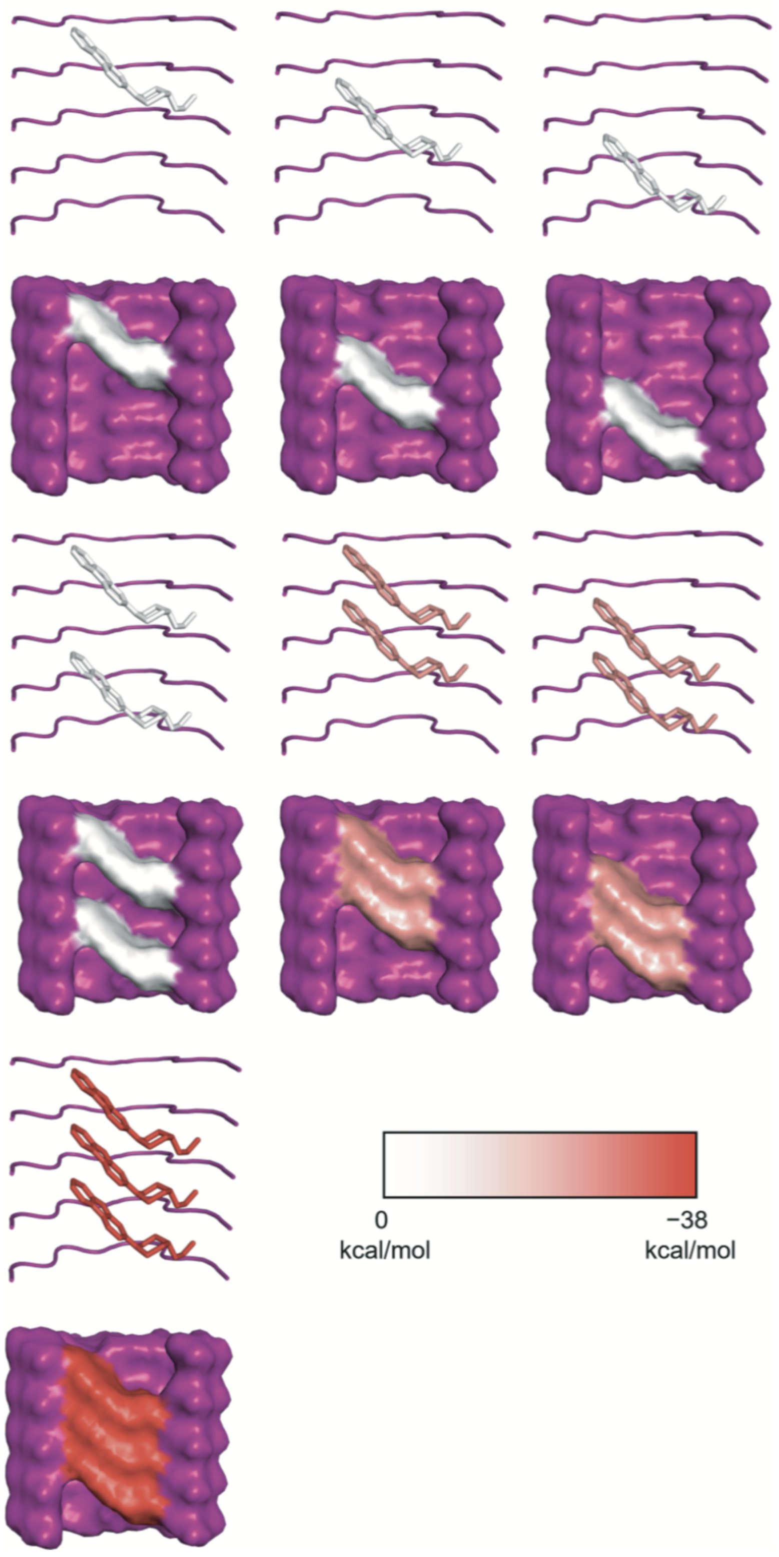


**Fig. S11. Energetics of GTP-1 binding to the amyloid filament.**
The ligands in the binding pocket (purple) with their energy represented along a color bar from 0 kcal/mol (blue) to −38 kcal/mol (red), with more negative energies being more favorable.

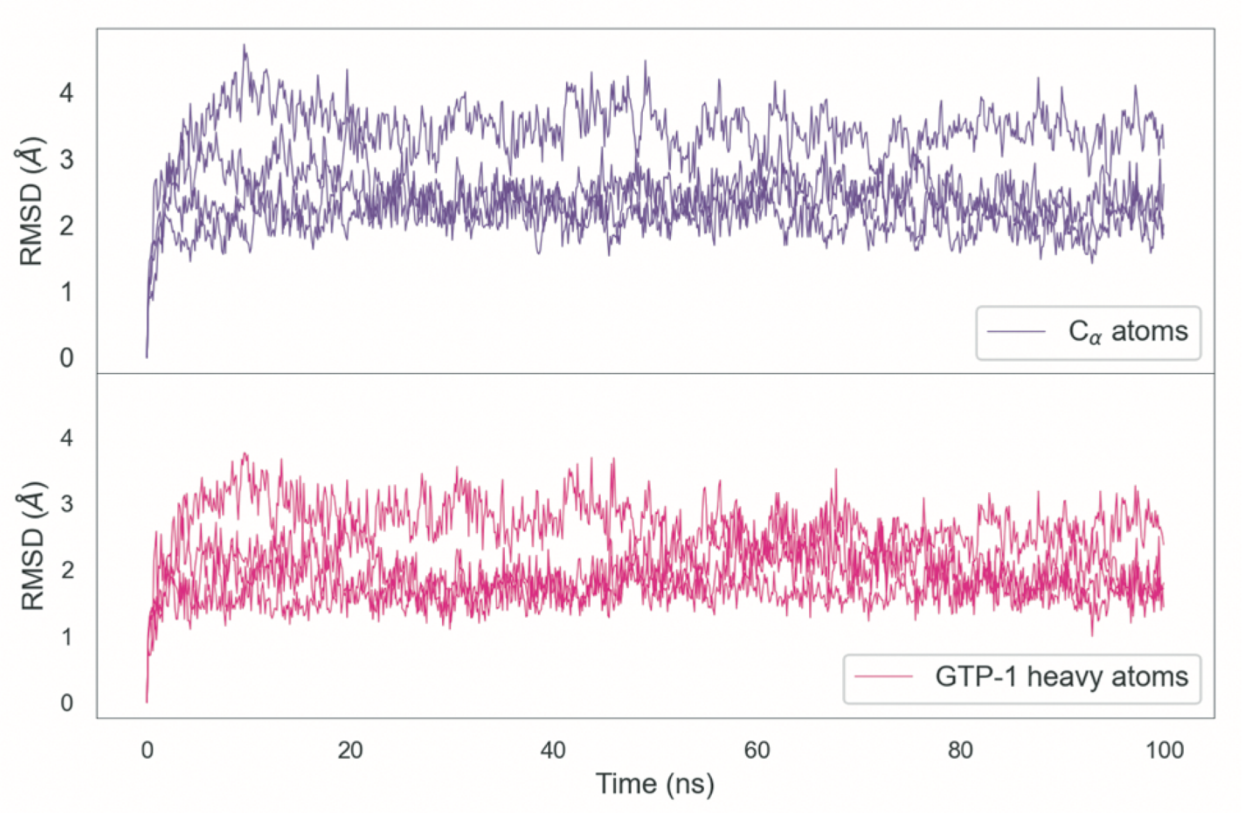


**Fig. S12. RMSDs of MD trajectories of tau PHF:GTP-1.**
RMSD of the $C_\alpha$ atoms of the tau PHF (top: purple) and the GTP-1 heavy atoms (bottom: pink) throughout a 100 ns MD simulation, showing the stability of the GTP-1 binding pose.

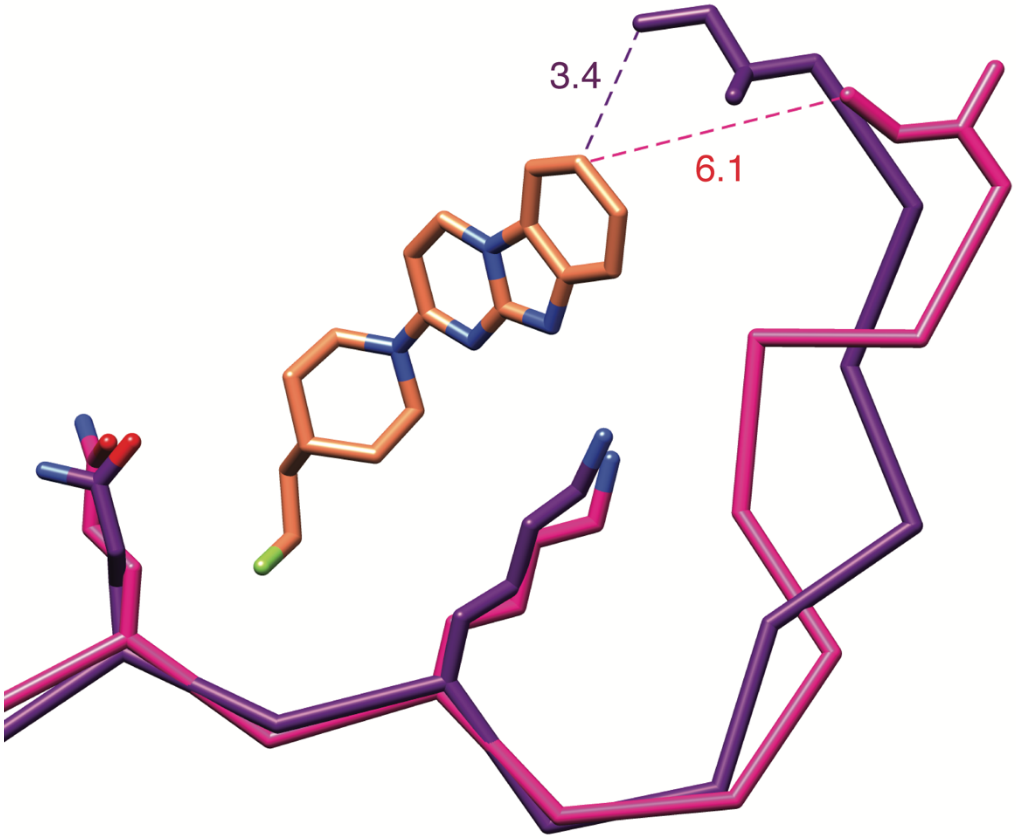


**Fig. S13. Comparison of the GTP-1 binding pocket in AD and CTE filaments.**
Residues 351–360 in AD (purple) and CTE (pink) filament structures, with the rotamer of Lys353 matching that in our AD + GTP-1 structure. The change in concavity of the pocket would move Ile360 away from GTP-1 and would prevent a productive apolar interaction with C7 on the GTP-1 heterocycle.

**Table S1. Cryo-EM data collection and refinement statistics.**

| | **PHF + GTP-1** |
|---|---|
| Microscope and camera | Titan Krios, K3 |
| Magnification | 105,000 |
| Voltage (kV) | 300 |
| Electron exposure ($e^{-}/Å^2$) | 46 |
| Dose rate ($e^{-}$/physical pixel/sec) | 16 |
| Exposure per frame (sec) | 0.024 |
| Defocus range (μm) | −0.8 to −1.8 |
| Physical pixel size (Å) | 0.834 |
| Movies collected | 15,160 |
| Box size (pixels) | 288 |
| Interbox distance (Å) | 28 |
| Initial segments extracted | 380,428 |
| Final segments | 30,199 |
| Resolution (Å) | 2.7 |
| B-factor ($Å^2$) | −42.9 |
| Helical rise (Å) | 2.37 |
| Helical twist (°) | 179.45 |

**Table S2. Refinement and model statistics of tau PHF: GTP-1**

| | Tau PHF:GTP-1 |
|---|---|
| **Model Composition** | |
| Non-hydrogen atoms | 5,810 |
| Protein residues | 730 |
| Ligands | 10 |
| **R.M.S. Deviations** | |
| Bond Lengths (Å) | 0.004 |
| Bond angles (°) | 0.988 |
| **Validation** | |
| Molprobity score | 2.30 |
| Clashscore | 6.91 |
| Rotamer outliers (%) | 3.1 |
| Cβ outliers (%) | 4.35 |
| **Ramachandran Plot** | |
| Favored (%) | 90.14 |
| Allowed (%) | 9.86 |
| Outliers (%) | 0 |
| PDB accession code | – |
| EMDB accession code | – |

**Table S3. Single point DFT calculations and surface area calculations of GTP-1 with tau.**

| Species | Electronic Energy (Hartrees) | $\Delta E_{binding}$ (kcal/mol) | $\Delta\Delta E_{binding}$ (kcal/mol) | Surface Area (Å$^2$) | ΔSurface Area (Å$^2$) |
|---|---|---|---|---|---|
| Apo | -18,371.533 | N/A | N/A | 3,743.6 | N/A |
| Optimized GTP-1 | -977.489 | N/A | N/A | N/A | N/A |
| Modeled GTP-1 | -977.483 | N/A | N/A | 531.3 | N/A |
| 2 Modeled GTP-1 | -1,954.997 | N/A | N/A | 768.2 | N/A |
| GTP-1 (top) | -19,349.073 | −35.5 | −0.3 | 3,472.4 | −1.2 |
| GTP-1 (middle) | -19,349.074 | −35.8 | −0.6 | 3,473.3 | −0.3 |
| GTP-1 (bottom) | -19,349.073 | −35.2 | 0 | 3,742.7 | −0.9 |
| 2 GTP-1 (top) | -20,326.644 | −90.6 | −19.6 | 3,658.5 | −85.1 |
| 2 GTP-1 (bottom) | -20,326.644 | −90.3 | −19.3 | 3,658.0 | −85.6 |
| 2 GTP-1 (gap) | -20,326.613 | −71.2 | −0.2 | 3,741.4 | −2.2 |
| 3 GTP-1 | -21,304.215 | −145.6 | −38.8 | 3,573.3 | −170.3 |
